# De novo whole-genome assembly and annotation of a high-quality coffee variety from the primary origin of coffee, *Coffea arabica* var. Geisha

**DOI:** 10.1101/2024.06.21.600137

**Authors:** Juan F. Medrano, Dario Cantu, Andrea Minio, Christian Dreischer, Theodore Gibbons, Jason Chin, Shiyu Chen, Allen Van Deynze, Amanda M Hulse-Kemp

## Abstract

Geisha coffee is recognized for its unique aromas and flavors and accordingly, has achieved the highest prices in the specialty coffee markets. We report the development of a chromosome-level, well-annotated, genome assembly of *Coffea arabica* var. Geisha, considered an Ethiopian landrace thatrepresents germplasm from the Ethiopian center of origin of coffee. We used a hybrid *de novo* assembly approach combining two long-reads single molecule sequencing technologies, Oxford Nanopore and Pacific Biosciences, together with scaffolding with Hi-C libraries. The final assembly is 1.03GB in size with BUSCO assessment of the assembly completeness of 97.7% of single-copy orthologs clusters. RNAseq and IsoSeq data were used as transcriptional experimental evidence for annotation and gene prediction revealing the presence of 47,062 gene loci encompassing 53,273 protein-coding transcripts. Comparison of the assembly to the progenitor subgenomes, separated the set of chromosome sequences inherited from *C. canephora* from those of *C. eugenioides.*, Corresponding orthologs between Geisha and Red Bourbon had a 99.67% median identity, higher than what we observe with the progenitor assemblies (median 97.28%). Both, Geisha and Red Bourbon contain an inversion on Chromosome 10 relative to the pseudomolecules of the genetic material inherited from the two progenitors that must have happened before the separation in the geographical migration of the two varieties. Lending support of a single allopolyploidization event that gave origin to *C. arabica* after the hybridization event with the two progenitor lines. Broadening the availability of high-quality genome assemblies of *Coffea arabica* varieties, paves the way for understanding the evolution and domestication of coffee, as well as the genetic basis and environmental interactions of why a variety like Geisha is capable of producing beans with such exceptional and unique high-quality.

## Introduction

Coffee is one of the most popular beverages worldwide, with an estimated consumption of 400 billion cups per year (Sacks et al 2019). While there are over 124 *Coffea* species (Davis et al 2011), the bulk of the coffee consumption comes from two species, *Coffea canephora* (Robusta) and *Coffea arabica* (Arabica). Arabica represents 58% of world coffee production and the remaining comes from Robusta (ICO, 2023). Arabica coffee has been the preferred coffee because of its refined taste, with a rich and well-balanced flavor profile (DeMatta et al, 2007).

*C. arabica*, has an allotetraploid genome with n=22 chromosomes derived from the hybridization of its maternal diploid progenitor species *C. eugenioides* (E genome) and its paternal progenitor, *C. canephora* (C genome), or robusta coffee, from a single allopolyploidization event about 100,000 years ago (Lashermes et al., 2016). In contrast to the other Coffea species, *C. arabica* is the only species that is largely autopollinated, with about 10% natural cross-pollination (Carvalho 1985). The center of origin and diversity of *C. arabica* is considered to be southwestern Ethiopia and southern Sudan (Lashermes et al., 2016, Montagnon et al., 2021). The original germplasm of the cultivated Arabica varieties outside Ethiopia transited from the center of origin to Yemen (Montagnon et al., 2021) from where it spread worldwide, giving origin to two similar germplasms, Typica and Bourbon, that later arrived in the Americas in the 17th and 18th centuries (Scalabrini et al. 2020). Currently, two high quality coffee genome assemblies have been produced, IGA-CARA 2.4 from the Red Bourbon variety (Scalabrini 2024) and CARA 1.0 from the variety Caturra that is derived from a single dominant mutation from Bourbon (Krug et al., 1949). Assemblies of the two progenitor species, *C. canephora* (Denoeud et al 2014) and *C. eugenioides* (NC_040035.1), are also available. Figure 1 shows a brief summary of the geographical origins of the current *C. arabica* varieties, emphasizing the unique origins of the variety, Geisha for which we report on its genome.

**Figure 1.**
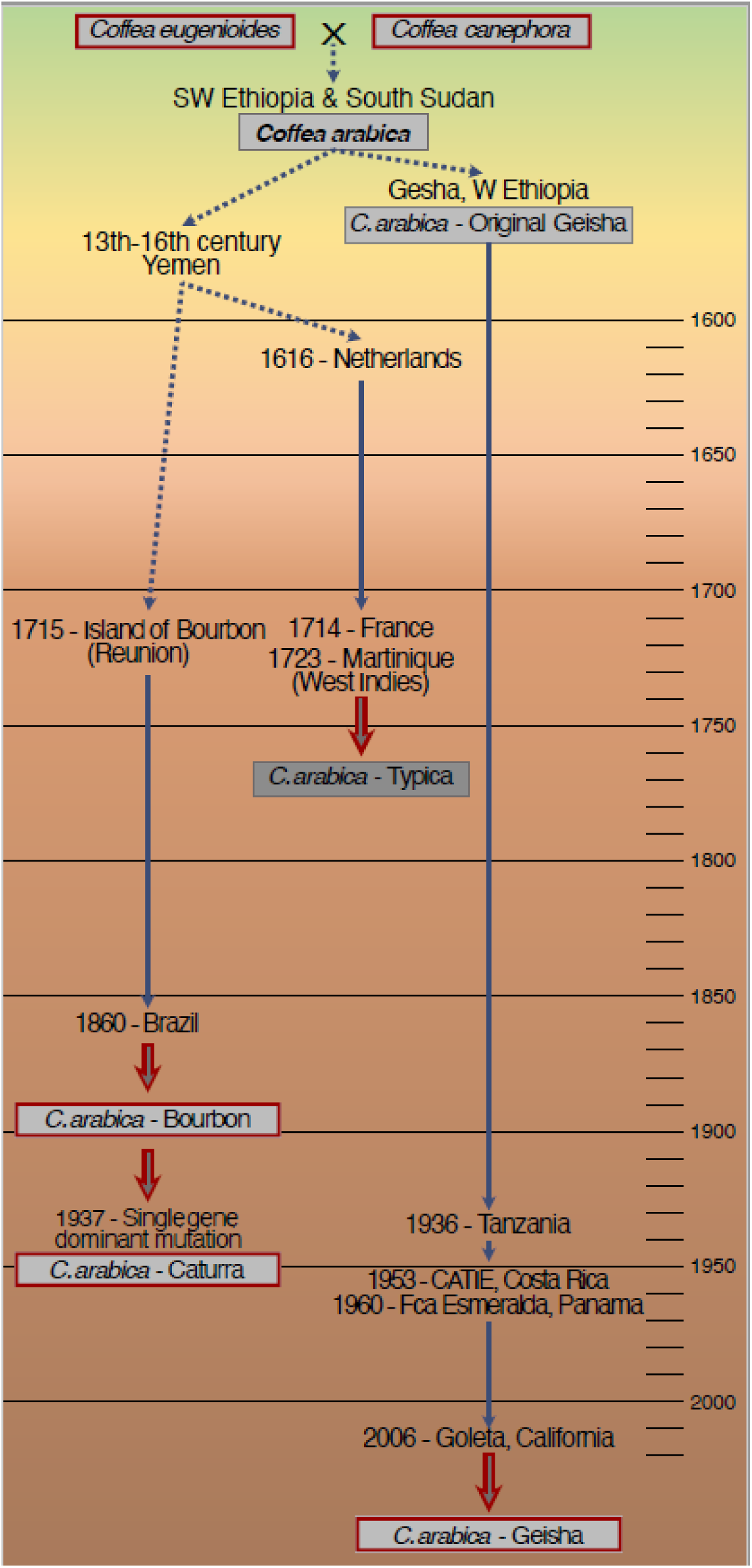
Time-line of origin and migrations of *Coffea arabica* varieties: *C. arabica* originated from the hybridization of the two diploid progenitor species *C. eugenioides* and *C. canephora* and a single allopolyploidization event. From the center of origin, one *C. arabica* variety, Bourbon, was derived from the early migration of *C. arabica* germplasm through Yemen*. C. caturra* later was derived from Bourbon by a single mutation event*. C. arabica,* Geisha, on the other hand, has a distinct direct connection to the center of origin of coffee and is considered a *C. arabica* landrace, as it travelled only in recent times from Ethiopia to California. (Species and varieties for which a genome assembly is available are highlighted with red boxes.) CATIE=Tropical Agricultural Research and Higher Education Center.

We developed a chromosome-level, well-annotated, genome assembly of *Coffea arabica* var. Geisha, considered an Ethiopian landrace, representing germplasm from the Ethiopian center of origin of coffee. The Geisha variety originated in the forests of the Gesha region in Western Ethiopia (Figure 1) and was collected around 1936 by Captain Richard Whalley, a British Consul for the Bench Maji region. Seeds were first sent to Kenya and then to the Lyamungu Research Station in Tanzania. From there, seeds were brought in 1953 to the Center for Tropical Agricultural Research and Education (CATIE) in Costa Rica (Boot, 2013). In the mid-1960’s, Geisha was planted from the CATIE germplasm in a small number of coffee farms in Boquete, Panama (Price Peterson, personal communication, Boot 2013). DNA analysis confirmed the origin of the CATIE Geisha as the original Geisha tree preserved in Tanzania (WCR 2016). Additionally, a study comparing Ethiopian forest Geisha type samples with Panamanian Geisha plants confirmed the high likelihood of the origin of the Panamanian Geisha from the Ethiopian forest (Krishnan, 2014). Geisha coffee gained prominence in 2005 for its unique, excellent quality when Hacienda Esmeralda, owned by the Peterson Family in Boquete, Panama, won the “Best of Panama Competition”, as well as setting a world record for highest auction price per pound of coffee at the time (WCR 2016). In 2006, a handful of Geisha coffee seeds were sent from Panama by Price Peterson to Jay Ruskey at Goodland Organics in Goleta, California. This introduction laid the foundation for the Geisha variety plants currently established at Goodland Organics and in Southern California, from where we collected plant material to develop our chromosome-level genome assembly (UCD 1.0).

In this study, we utilized long-read sequencing based on Oxford Nanopore Technologies (ONT, Oxford, England) and Pacific Biosciences (PacBio, Menlo Park, USA), together with scaffolding with Dovetail (Scotts Valley, USA) genome-wide chromatin conformation capture technologies, to develop a chromosome-level, well annotated, genome assembly of *C. arabica* variety Geisha. This paves the way for understanding the evolution and domestication of coffee, as well as the genetic basis of why a variety like Geisha is capable of producing beans with such exceptional and unique high-quality, and how environmental interactions like altitude and changing climate parameters can affect quality and yield in coffee.

## Materials and Methods

### Plant material collection and extraction of nucleic acid

Young leaf tissues were collected from coffee plant UCG-17 of variety ‘Geisha’ (*Coffea arabica*) at Goodland Organics in Goleta, California. For long-read sequencing, leaves from the second youngest node were processed using a modified CTAB extraction protocol as described in Stoffel et al. (2015) to produce high molecular weight (HMW) DNA. Second node leaves were also utilized for DNA extraction using the Qiagen DNeasy Plant Kit for short read sequencing. DNA concentration was measured using QuBit (Thermo Fisher Scientific, Waltham USA) and fragment size and integrity was evaluated with an Agilent 2100 Bioanalyzer (Agilent Technologies, Inc., Santa Clara, USA).

For transcriptome sequencing, we collected samples of different organs and tissue types in RNAlater (Thermo Fisher Scientific, Waltham, USA) at different developmental stages (Table S1), from Geisha plants at the same Goleta, California location. For RNASeq, each tissue was extracted using Qiagen RNeasy Plant Kits. For IsoSeq, RNA was extracted using the CTAB method as reported in Blanco-Ulate et al. (2013). The integrity of RNA samples was evaluated on an Agilent 2100 Bioanalyzer (Agilent Technologies, LLC).

### Genome size estimation with flow cytometry

For genome size estimation, a ∼50 mg sample of coffee leaves from the Geisha plant UCG-17 was collected in Goleta, California and shipped between moist paper towels on cooling packs for estimation of genome size using flow cytometry. Intact nuclei preparation was performed at Benaroya Research Institute, Seattle, WA and nuclear content was estimated by taking an average of four repeated estimates. The resulting estimate of genome size was 2.22±0.030 pg/2C or 1.09 Gb haploid genome size.

### Genomic libraries preparation and sequencing

A whole genome sequencing Illumina library with an average insert size of 350 bp was sequenced on the Illumina HiSeq X machine producing a total of 37X genome coverage with 150 bp paired end reads. Quality of sequencing was assessed with FASTQC software. Reads were processed to remove adapters and trim low quality bases using CLC Genomics Workbench (Qiagen, Valencia, USA).

Pacific Biosciences SMRT sequencing libraries were constructed according to manufacturer recommendations at the UC Davis Genome Center. Fragments >10kb were size-selected for sequencing via BluePippin (Sage Science, LLC). The library was sequenced using 101 SMRT cells of the P6-C4 technology on the RS II. A total of 95.3 Gigabases (Gb) of sequence was produced equivalent to 79.4X genome coverage. N50 of reads ranged from 16-21Kb. Cleaned reads were utilized for genome assembly with FALCON-UNZIP followed by two rounds of Quiver for base correction, to obtain primary contigs and secondary haplotigs. A final round of polishing was performed using PILON software (Walker et al, 2014) and whole genome sequencing short-read data to correct for SNPs and small indels.

An Oxford Nanopore Technologies library was constructed with 250 Kb averaged size DNA fragments according to manufacturer instructions at the UC Davis Genome Center. The library was sequenced on two flow cells on a Promethion (Oxford Nanopore Technologies) which generated 150X genome coverage. The CANU v1.8 software (Koren et al, 2017) was used to produce an assembly with the ONT sequence. The resulting assembly was polished with the ONT raw reads using Racon software (Vaser et al., 2017), resulting in ONT assembly v0.5.

Chicago and HiC libraries (Dovetail) were produced from HMW DNA. The chicago library was sequenced and produced 115.0 Gbp of data, corresponding to a depth of genome coverage of 88.5x X-Fold. HiC library sequencing produced 84.5 Gbp of reads, corresponding to 65x X-Fold coverage. The standardized Dovetail informatics pipeline (HiRise) was utilized with the FALCON-UNZIP polished primary contigs fasta files as input. Briefly, the pipeline first processes the Chicago sequencing data and corrects short range distance information, then utilizes the HiC sequencing information for further scaffolding of long-range information. A number of breaks/joins were investigated manually to confirm the software functionality. Plots showing the resulting HiC linkages were also investigated manually. This process utilizing the Chicago and HiC libraries was repeated utilizing the v0.5 ONT assembly, after which this scaffolded version of the ONT assembly was polished with an additional round of Racon (Vaser et al., 2017) using the Pacific Biosciences raw reads to produce v1.0 ONT Dovetail assembly. Plots showing the resulting HiC linkages were investigated manually. Standard assembly quality statistics were evaluated for each assembly version, including gaps versus number contigs and scaffolds.

### Assembly comparisons and pseudomolecule generation

The PacBio Dovetail assembly was aligned to the v1.0 ONT Dovetail assembly using RaGOO (Alonge et al., 2019). Scaffolds in the PacBio Dovetail assembly were further scaffolded according to the alignment with the v1.0 ONT Dovetail assembly to produce a final draft assembly. This final draft assembly was aligned with MUMmer software (nucmer algorithm) (Kurtz et al., 2004) to determine relationships with the progenitor genome assemblies available at NCBI - *C. canephora* (GCA_900059795.1) and *C. eugenioides* (GCA_003713205.1). Global alignments were visualized with minimum lengths of 5,000 bases and both 98% and 99% identity between the sequences. Pseudomolecule pair alignments that dropped off by increasing the percent identity to 99% represent the scaffold corresponding to the opposite subgenome homeolog. The correlation of this identity was used to assign and name the pseudomolecules to each subgenome. Chromosome pairs were numbered following the *C. canephora* genome (Dereeper et al., 2013), with ‘c’ or ‘e’ indicating the subgenome of inheritance from *C. canephora* or *C. eugenioides*, respectively. For example, Chr01e refers to ortholog to Chromosome 1 of the *C. eugenioides* genome while Chr01c refers to ortholog of Chromosome 1 of the *C. canephora* genome. The resulting assembly arranged in named pseudomolecules or chromosomes is designated the final version as Coffee (*Coffea arabica*) genome UCD v1.0 for variety Geisha. The completeness of the final genome assembly was assessed by search of highly conserved single-copy orthologs with BUSCO V5.1 (Simao et al. 2015) using Eudicots odb10 database of gene models.

### Genome Annotation

Repetitive sequences were identified in the final draft assembly using RepeatModeler 2.0 (Smit and Hubley 2019), to identify repeat families present specifically in the coffee genome. A custom coffee repeat library was produced and then used as input with the final genome assembly with RepeatMasker 4.1.0 (Smit et al. 2013) to identify repeat positions in the genome and to generate a masked version of the genome sequence. This masked version was used as the input for the annotation procedure that followed.

High-quality IsoSeq data were used to produce high-quality gene models for training gene predictors in PASA v.2.4.1 (Haas 2003) along with transcript evidence obtained from RNAseq data aligned on the genome using HISAT2 2.1.0 (Kim et al. 2015) and by performing transcriptome assemblies using Stringtie v.2.0 (Pertea et al. 2015) and Trinity v.2.8.5 (Grabherr et al. 2011). Public databases (Swiss-Prot and UniProt Nov. 11, 2020), transcriptome assemblies, and the Iso-Seq data described above were used as transcript and protein evidence. They were mapped on the genome using PASA v.2.4.1 (Haas 2003), and Exonerate v.2.4.0 (Slater and Birney 2005). Ab initio predictions were generated using Augustus v.3.3.3 (Stanke et al. 2006), GeneMark v.4 (Lomsadze 2005), and GlimmerHMM 3.0.4. EvidenceModeler v.1.1.1 (Haas et al. 2008) and PASA used these predictions to generate consensus gene models. Models showing untranslated regions (UTRs) or introns longer than 25 kb were removed, as well as models encoding for proteins with no orthologue correspondence (50% of identity and coverage) with other known plant proteins in the RefSeq Database (January 17, 2017). The final functional annotation was produced combining Diamond v.2.0.4 (Buchfink et al. 2014) hits against the UniProt protein database and InterProScan v.5.40-77.0 (Jones et al. 2014) outputs using Blast2GO v.4.1.9 (Gotz et al. 2008).

### Comparison of genome assembly and synteny analysis

Putative pseudogenes in *C. arabica* var. Geisha genome were identified by reciprocal mapping of the annotated CDS sequences using BLAT v.36×2 (Kent 2002). Alignments with an identity greater than 50% and a coverage of 80% of the CDS sequence were considered to allow flexibility in terms of sequence identity while ensuring the match would encompass significantly conserved genomic regions. Only those matches not overlapping any other annotated gene locus and encoding for an incomplete ORF (ex. premature stop codon, lack of methionine at start, etc.) were considered putative pseudogenes.

Relationships between ortholog protein-coding genes of the two subgenomes and the parental species *C. canephora* (NCBI assembly and annotation GCA_900059795.1) and *C. eugenioides* (NCBI assembly and annotation GCF_003713205.1) were established searching for homolog proteins with BLASTp v.2.7.1+ (Altschul et al. 1990), using a minimum e-value 0.001 and collecting only the best 100 hits. Synteny between the progenitor species with the corresponding subgenomes and between the two subgenomes was performed with MCScanX v.11.Nov.2013 (Wang et al. 2012). To relate homologous loci across different genomes, coding genes were associated using the hits from protein BLAST searches; pseudogenes were associated with annotated coding genes. Syntenic blocks were defined with a minimum of 15 genes per block and a maximum stretch of 10 genes missing per block.

The assembly structure of *C. arabica* variety Geisha was compared with the assembly of *C. arabica* variety Red Bourbon from (Scalabrin et al. 2024) (IGA_Cara_2.4, NCBI accession GCA_030873655.1). Whole genome sequence alignments were produced between each pair of genomes and subgenomes using Minimap2 v. 2.26-r1175 (parameters: “-k19 -w19 - U50,500 -A1 -O6,26 -s200”) (Li et al. 2018). To compare the sequence identity across multiple assemblies, *C. arabica* variety Geisha and *C. arabica* variety Red Bourbon genomes sequences were sliced into 25Kbp windows using makewindows (parameters: “-w 25000”) and getfasta tools from the BEDtools suite v2.24.0 (Quinlan and Hall, 2010). The sequences of the 25Kbp windows were then mapped against all other subgenomes pseudomolecules and progenitor genome assemblies (*C. canephora* and *C. eugenioides*) using Minimap2 v. 2.26-r1175 (parameters: “-x map-hifi”) (Li et al. 2018). The coverage and identity for each hit were calculated. For each genomic window in each comparison, the hit showing the highest number of matching nucleotides was selected as the best one.

## Results and Discussion

### *Coffea arabica* genome size estimates

Coffea species have been estimated to have a wide range of genome sizes, from the smallest reported diploid genome of 0.95±0.13 pg in *C. racemosa* (Cros et al. 1995) to the largest in the genus ascribed to the *C. arabica*. This is consistent with *C. arabica* being an allotetraploid and, thus, having inherited two subgenomes by a hybridization event from two diploid progenitors, *C. canephora* and *C. eugenioides.* Both of the diploid progenitors of *C. arabica* appear to have fairly consistent genome size estimates using flow cytometry. *C. eugenioides* has a reported 2C of ∼1.36 pg and *C. canephora* a slightly larger 2C of ∼1.46 pg (Table S2). However, the estimates for *C. arabica* not only have been reported to be more variable (Table S2) than the progenitors, but also to be consistently smaller than their combined size of ∼2.82 pg.

Our flow cytometry analysis of *C. arabica* variety Geisha genome size reported a 2C value of 2.22±0.030 pg. This estimate is marginally smaller than estimates previously reported for other *C. arabica* varieties (2.30pg∼2.72 pg) but within the range of variability previously observed due to the occurrence of “intraspecies polymorphism”, the estimating technique used and the application of the methods to a tetraploid genome (Cros et al. 1994, Clarindo et al. 2013). From our flow cytometry estimate, the 1C nuclear genome size of *C. arabica* var. Geisha should approximate 1.085 Gbp (assuming 1pg = 978 Mb, Dolezel et al. 2003).

### Assembly of Geisha genome

The genome of *C. arabica* variety Geisha (UCDv1.0) was reconstructed using a hybrid *de novo* assembly approach combining two long-reads single molecule sequencing technologies (Pacific Biosciences SMRT technology and Oxford Nanopore ONT technology), together with Dovetail proximity-ligation sequencing technologies for scaffolding both assemblies (Table 1, Figure S1).

**Table 1.** Genome assembly and annotation statistics.

| **Assembly** | ***Geisha -UCD1.0*** |  |
| --- | --- | --- |
| **Assembly length (Kbp)** | 1,025,607 |  |
| **Chromosomes** | 22 |  |
| **Unplaced sequences** | 214 |  |
| **%GC** | 37 |  |
| **Number of gaps** | 1,876 |  |
| **Total gap length (Kbp)** | 223.6 |  |
| **Median scaffold (Kbp)** | 134.9 |  |
| **Maximum scaffold (Kbp)** | 69,315.2 |  |
| **Scaffold N50 (Kbp)** | 43,715.0 |  |
| **Scaffold L50 (#)** | 10 |  |
|  | ***ONT based*** | ***PacBio Based*** |
| **Scaffold N50 (Mbp)** | 44.49 | 42.54 |
| **Scaffold L50 (#)** | 12 | 10 |
| **Total contigs** | 2,112 |  |
| **Contig N50 (Kbp)** | 1,107.8 |  |
| **Contig L50 (#)** | 234 |  |
| **Annotation** |  |  |
| **Number of gene loci** | 47,062<br>(8,617 pseudogenes) |  |
| **Number of proteins** | 53,273 |  |
| **Number of complete BUSCO** | 2,272 |  |
| **Number of missing BUSCO** | 30 |  |
| **Complete BUSCO (%)** | 97.7 |  |
| **Repeat (%)** | 60.6 |  |

(V5.1, Eudicots odb10 database, 2,326 BUSCOs)

Long reads datasets were first assembled separately. PacBio SMRT sequencing produced approximately 87.8X coverage based on the 1.09 Gbp genome size estimated by flow cytometry. PB long reads were assembled using the FALCON-Unzip pipeline (Chin et al. 2016) producing an assembly containing 1,316 primary contigs comprising 1.025 Gb total bases (94.42% estimated size) with N50 of 1.85 Mb. The primary assembly size suggested the assembly reported a non-redundant representation of one single haplotype for both subgenomes C and E.

Oxford Nanopore Technologies (ONT) long read sequencing generated ∼150X-fold coverage of the genome and was assembled using CANU (Koren et al. 2017) which produced 1,685 contigs comprising 1.18 Gbp (109% estimated size) with N50 of 4.10 Mb. The CANU assembly procedure, unlike FALCON-Unzip, is not haplotype-aware, and the increased assembly size with respect to the estimated flow cytometry can be attributed to the lack of phasing and compressing of the alternative alleles and causing the co-presence of alternative haplotype sequences.

After polishing, both sets of assembled contigs underwent two-stage scaffolding using two different Dovetail Genomics proximity-ligation sequencing technologies. The first scaffolding stage involved the use of a Chicago library, the second stage involved a HiC library. Comparing the overall assembly statistics after scaffolding of the PB and ONT assemblies (Table 1), showed that the ONT assembly was approximately 1.04 times more contiguous based on scaffold N50 of 44.48 Mb in ONT versus 42.54 Mb in PB.

Initial BUSCO (Simao et al. 2015) (V3 with Eudicots odb9 database with 2121 gene models) assessment showed that the PB assembly located 2,058 (97.1%) complete genes while the unpolished ONT assembly resulted in 1,124 (53.0%) complete BUSCOs. After several polishing steps with both Nanopore ONT and PacBio SMRT long reads, it was possible to locate as complete loci 94.7% (2,008) of the BUSCOs on the ONT-based assembly. The use of long-read for polishing the ONT-based assembly sequences increased the quality at base level, however, improvement did not reach the PacBio-based assembly quality.

Therefore the two assemblies were then combined using RaGOO (Alonge et al., 2019), using the higher contiguity of the ONT-based assembly to guide the scaffolding of the higher base-quality PacBio assembly sequences (Figure S1) and further reduce the number of scaffolds to 236.

### Pseudomolecule creation and gene annotation

To generate the final assembly, we used a very effective approach combining the superior contiguity of the ONT v1.0 Dovetail assembly and the base quality of the PB Dovetail assembly. RaGOO and MUMmer software were utilized to generate alignments between these two assemblies, then order and orient the PB Dovetail assembly with the ONT v1.0 Dovetail assembly to produce the final genome assembly - Coffee (*Coffea arabica*) genome UCD v1.0. The final assembly contains 1,025,606,995 bases which is 94.5 % of the estimated genome size of 1.09 Gb by flow cytometry of the same plant. The assembly has 96.8 % of the sequence bases in 22 pseudomolecules corresponding to the 22 chromosomes. The remaining 3.2% of the sequence is in only 214 remaining scaffolds, all greater than 20 kb (Table 1).

BUSCO assessment of the completeness of the final assembly (Table 1) presented a representation of 2,272 out of the 2,326 single-copy orthologs clusters (97.7%) in the eudicotyledons database (odb10). This figure is comparable with the Caturra and Red Bourbon genomes (95.8 and 99.7% respectively), confirming the high completeness of the assembly and comprehensive representation of the *C. arabica* var. Geisha gene space.

The analysis of repetitive content in the Geisha genome classified 60.6 % of the assembly as repetitive. This fraction is consistent with the other Coffea genomes (*C. eugenioides* 61.5%, *C. arabica* var. Caturra 62.5 %, *C. arabica* var. Red Bourbon 59.2 %), with the exception of *C. canephora* where a figure of 44.8 % suggests an incomplete representation of the repetitive content, likely due to technical limitations of the assembly.

RNAseq and IsoSeq data were used as transcriptional experimental evidence for annotation and gene prediction of the Geisha genome. The PacBio full-length cDNA isoforms were first combined with the mRNA short read assemblies, first to guide the training of *ab initio* gene predictors and then to polish the final predictions to obtain alternative splicing events (Figure S2). The annotation of the Geisha genome revealed the presence of 47,062 gene loci encompassing 53,273 protein-coding transcripts (Table 1). Comparing the gene content of the two subgenomes we could identify 8,617 conserved loci that did not encode for a full ORF and have potentially become pseudogenes. The number of loci is consistent for an allotetraploid with the reported numbers from the progenitors’ genomes (25,574 in *C. canephora* and 33,619 in *C. eugenioides*) (Table S3).

### Comparison with progenitor species

By comparing the pseudomolecules of the UCD v1.0 assembly to the progenitor genomes available in NCBI, it was possible to separate the set of chromosome sequences inherited from *C. canephora* from the ones inherited from *C. eugenioides* (Figure 2). Selecting only the alignments with 99% identity between sequences shows a polarization of the hits of each homeolog with just one progenitor and the complete loss of alignment across the full length of the pseudochromosome to the sequence of the other progenitor (Fig 2A and Fig 2B). As observed for Red Bourbon (Scalabrin et al. 2024), for some chromosomes like chromosome 10, we observed a switch of the inherited genetic content between homeologs, with a portion of the pseudomolecule originating from the other progenitor species (Fig 2B).

**Figure 2.**
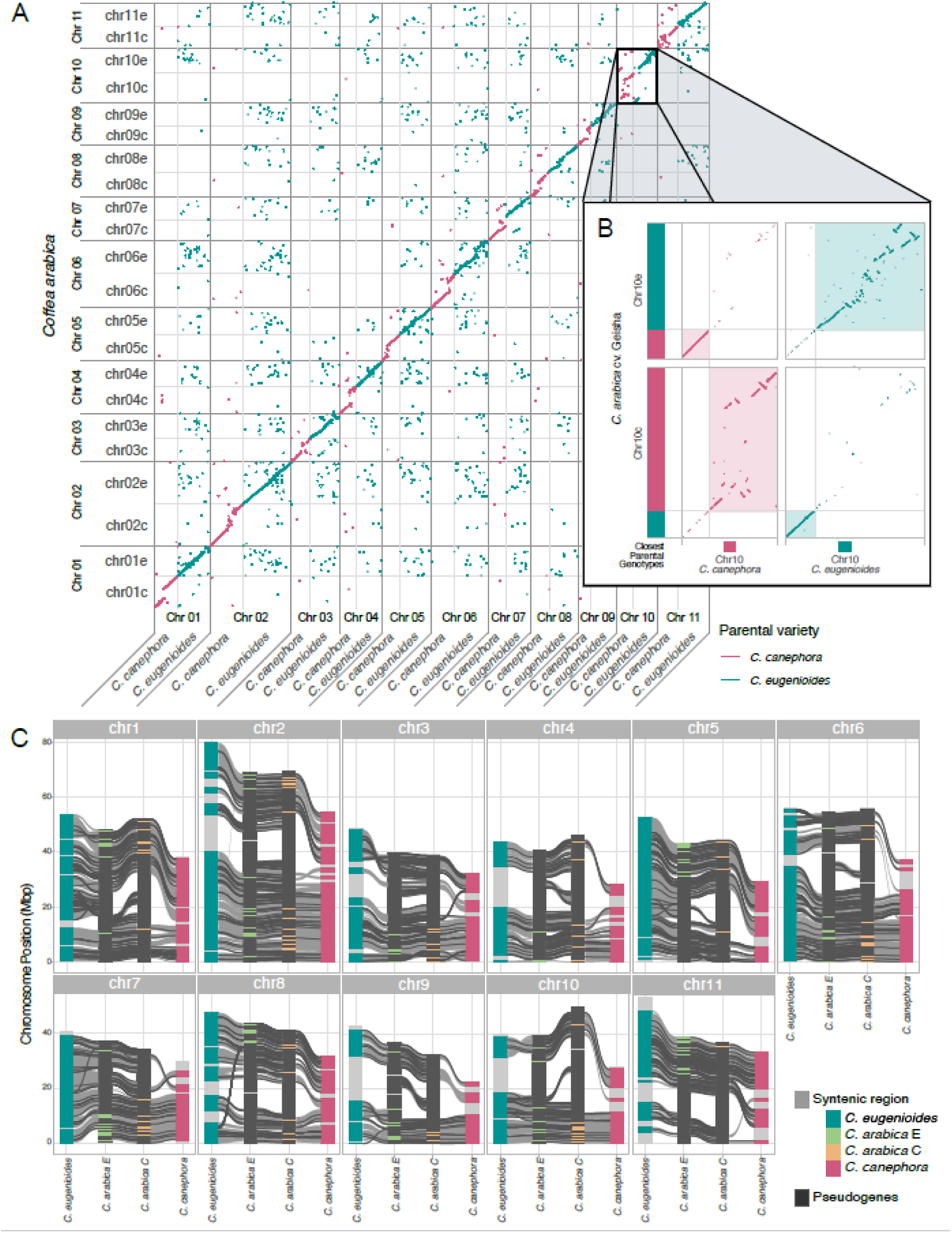
Comparison between *Coffea arabica* - Geisha and the progenitor species. **A)** Dot plot diagram comparing the Geisha pseudomolecules to the progenitor chromosome sequences to identify the origin of the subgenomes. Box highlights location of chromosome 10. **B)** Expanded view of the alignment of chromosome 10 showing the closest parental genotypes *C. canephora* and *C. eugenioides* for each segment of the pseudomolecules and the switch of inherited content between homeologs. **C)** Synteny comparison of the chromosome sequences of Geisha and the progenitors species. *C. arabica* E is orthologous to *C. eugenioides* and *C. arabica* C is orthologous to *C. canephora*. Grey connecting ribbons show syntenic blocks of conserved gene loci between pseudomolecules. Black lines on chromosomes indicate the location of putative non-coding pseudogene loci.

The chromosome length, sequence and structure of the pseudomolecules are closer between the homeologs within *C. arabica,* rather than each subassembly with the original progenitor genome (Fig 2C). *C. eugenioides* pseudomolecules were overall longer than the associated homeologs in *C. arabica*, confirming what was observed with the flow cytometry that genome size of *C. arabica* is expected to be smaller than the combined length of the progenitors (2.22pg vs ∼1.36pg + ∼1.46pg). All *C. canephora* pseudomolecules, on the contrary, show a reduced size when compared to the other assemblies, a situation contradicting the flow cytometry data that would suggest a genome size comparable if not larger than *C. eugenioides*, likely due to a lower assembly quality for the species when compared to the other progenitor.

The comparison of gene content confirms what was observed at the sequence level. Of the annotated loci in the Geisha genome, 89.5% result in syntenic collinear blocks between homeologs. The number drops to 79∼82% when comparing each subgenome set with the respective progenitor, for *C. canephora* and *C. ceugenioides*. Interestingly, for most of the chromosomes, we can observe a consistency across the intra-species and inter-species comparisons of which regions appear to be syntenically conserved and regions losing structural coherence (Fig 2C). Where we observe a loss of synteny between homeologs, we observe a loss of synteny at least with one of the progenitors. Moreover, those regions seem to accumulate and concentrate pseudogenes (Figure 2C), suggesting a higher variability or instability leading to the accumulation of deleterious variants for one of the two homeolog loci.

### Comparison to the Red Bourbon genome sequence

The genomes of Geisha and Red Bourbon varieties were assembled with similar procedures (Scalabrin et. al 2024), making them the most comparable from a technical perspective. The structure of all pseudomolecules of both subgenome assemblies C and E is remarkably more similar between the two varieties rather than what can be observed when comparing the subgenome assemblies with each of the respective progenitors (Figure 2A and Figures S3 and S4). At the nucleotide level, we observe a similar situation. Comparing Geisha and Red Bourbon, corresponding homeologs shows a 99.67% median identity (Figure 3A, Figure S5A), higher than what we observe with the progenitor assemblies (median 97.28%, Figure S5B). Cross-comparing the subgenomes E with subgenomes C shows a homogeneous picture, with a lower median identity of ∼92% regardless if the pair of subgenomes come from the same variety (Figure 5SC).

**Figure 3.**
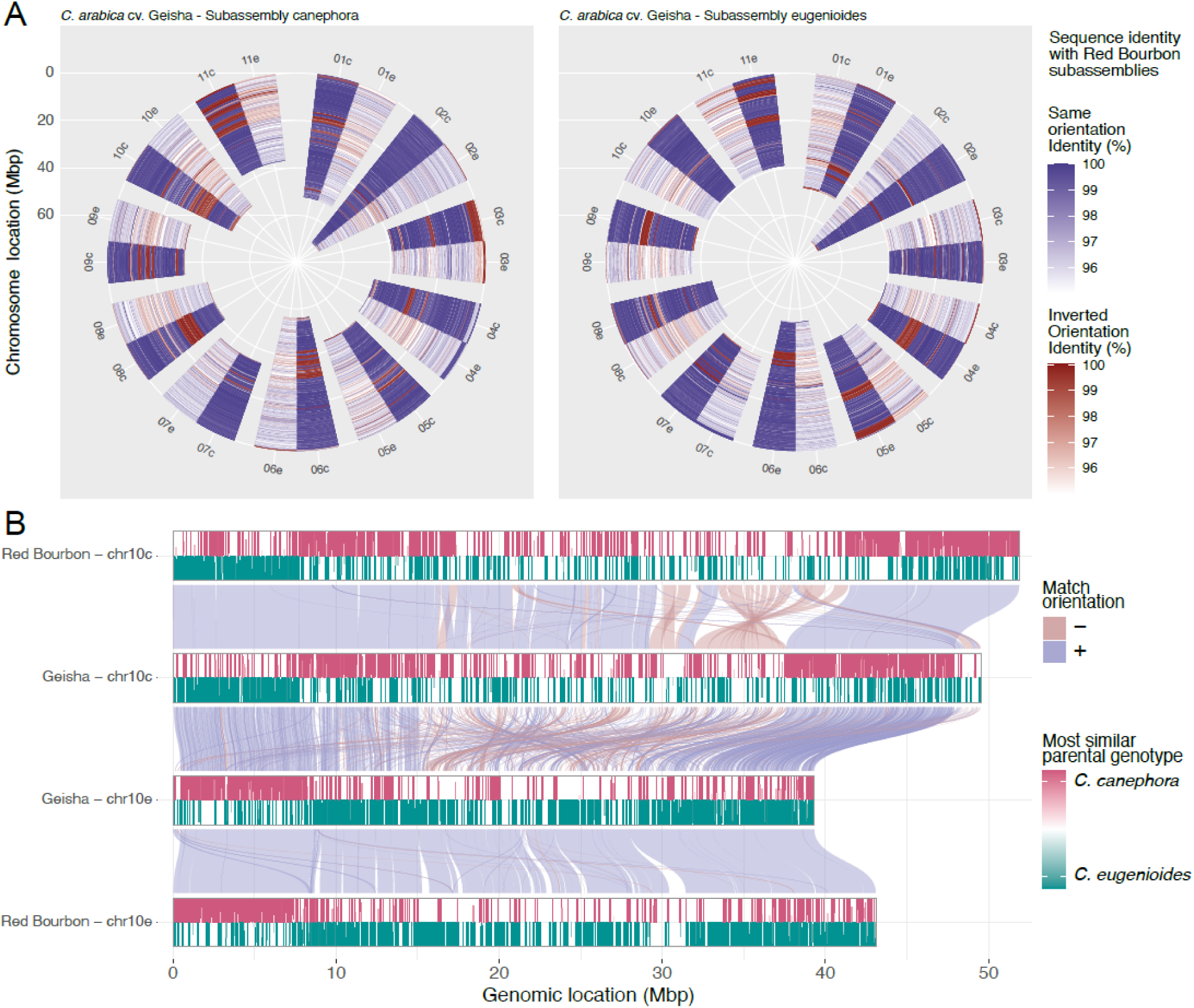
Comparison between the assemblies of *Coffea arabica* variety Geisha (UCD_1.0) and Red Bourbon (IGA_CARA_2.4). **A)** Polar diagram of the pseudomolecule identity levels between Geisha and Red Bourbon (Blue scale indicates sequence homology in the same orientation. Red indicated sequences in inverted orientation). Different levels from the center of the diagram correspond to positions across the length of the chromosomes. **B)** Aluvial plot showing homolog regions between chromosome 10 pseudomolecules of Geisha and Bourbon varieties. Chromosome bars show color-coded homology levels of pseudomolecule region with the progenitor species (red *C. canephora*, green *C. eugenioides*).

Despite the presence of translocation and inversion events in all homeolog pseudomolecule comparisons, the subassembly E appears to be slightly more conserved in both structure and nucleotide content between varieties and with the progenitor species than their *C. canephora* counterpart (Fig 2C, Fig S5).

Particularly interesting is the case of Chromosome 10, which contains an inversion in the pseudomolecules of the genetic material inherited from the two progenitors. These kinds of conditions are generally ascribed to errors in the assembly procedure, as polyploid genomes can cause the assembly algorithms to switch haplotypes (Chin et al. 2016). However, a similar situation has been observed in Chromosome 10 pseudomolecules in the independent assembly of Red Bourbon (Scalabrin et. al 2024). By comparing the two sets of Chromosome 10 homeologs (Figure 3B), we can confirm that both varieties bear a chromosomal exchange event between the two progenitor lines. Moreover, the identity in sequence and structure around the switch breakpoints, for both subgenomes of both varieties, suggests that they originated from the very same ancestral recombination event. It is also possible to observe the higher coherence in the structure and the sequence of the regions inherited from C. *eugenioides* than that inherited from *C. canephora*. While they are very similar in sequence content, they show a higher extent of structural rearrangements, and this happens in correspondence of the region that is most dissimilar between the progenitors lines (Figure 3B). This region appears to be missing or under-assembled in the *C. canephora* genome reconstruction (Figure 2C), although evolution of the *C. arabica* Chr10c after inheritance from its progenitor cannot be discounted.

It is impossible to trace the evolution of *C. arabica* from the genomic data of just a few reference genomes, however the distant domestication history between Geisha and Red Bourbon varieties allows us to trace them back to a common origin. The nucleotide identity and the concomitance of the recombination event in chromosome 10 suggests that both varieties derived from the same hybridization event between *C. canephora* and *C. eugenioides*. The evolution of the parental genetic material in chromosome 10 must have happened before the separation of the two varieties and the export of the *C. arabica* line that led to the Red Bourbon line from Ethiopia to Yemen to Reunion (Figure 1). This lends support to Lashermes (2016) conclusion of a single allopolyploidization event that gave origin to *C.* arabica after the hybridization event with the two progenitor lines and is consistent with Salojärvi et al. (2024). Evolution, breeding, and domestication caused the accumulation of the other structural variants we observe that differentiate the structure of the pseudomolecules of Geisha and Red Bourbon. Moreover, the plasticity of the genome appears to be very variable across the different homeologs and different chromosomes.

Comparing different varieties of *C. arabica*, we can assess the quality of genome assemblies and, at the same time, investigate the evolution of the species, impact of domestication events, and allow for precision breeding. Availability of more Coffea genomes would also provide a model to understand the mechanisms of allopolyploidization in plants, as it has been observed that when crossing *C. arabica* with a progenitor line hybrid it can produce an allohexaploid hybrid, Clarindo et al. (2013). Currently few resources are available, as only two other *C. arabica* assemblies are accessible (Caturra and Red Bourbon), both resulting from the same line of domestication and distribution (Figure 1). Therefore, it is important to broaden the availability of high quality genome assemblies of more Arabica varieties, the Arabica progenitor lines, and other wild Coffea species.

## Data availability

Sequencing data are accessible at the NCBI repository under the accession PRJNA1032043. The Whole Genome Shotgun project has been deposited at DDBJ/ENA/GenBank under the accession JBELAZ000000000. The version described in this paper is version JBELAZ010000000. Final assembly data are available in Zenodo under the DOI 10.5281/zenodo.1003885 and in Phytozome, genome ID 871, https://phytozome-next.jgi.doe.gov/info/Carabicageisha_1_0.

Pipeline/scripts are available on G3 Figshare 405138

## Acknowledgements

This work was funded by a gift from Suntory Global Innovation Center Limited, Tokyo, Japan. We thank Yoshi Tanaka (Suntory Global Innovation Center) for his support throughout the development of this project. Support also was provided by the UC Davis Department of Animal Science, UC Davis Seed Biotechnology Center, UC Davis Department of Viticulture and USDA/ARS, Raleigh, NC. We also gratefully acknowledge the work by David Goodstein and Shengqiang Shu from Phytozome for hosting and cross-checking the genome and annotation. We thank Jay Ruskey (Goodland Organics and Frinj Coffee) for making available the plant material described in this study and for his friendship and collaboration. We thank Alma Islas (UC Davis) for preparing the tissues and libraries for RNA analysis. We also gratefully acknowledge the support of Oanh Nguyen at the UC Davis Genome Center for all the sequencing work in this study.

---

**Figure S1: *C. arabica* cv. Geisha UCDv1.0 genome assembly pipeline.** The genome was reconstructed using a novel hybrid *de novo* assembly approach to combine two long-reads single molecule sequencing technologies (Pacific Biosciences SMRT technology and Oxford Nanopore ONT technology), together with Dovetail proximity-ligation sequencing technologies for scaffolding both assemblies. Assembly statistics are shown for the progression of the different steps of the assembly.

**Figure S2: Genome annotation pipeline.** Input for the annotation was a masked version of the genome. RNAseq and Iso-Seq data from 10 different Geisha coffee tissues (Table S1) served as external evidence for gene prediction. Mapped transcriptional data was consolidated to generate a training set for ab-initio predictors. Together with protein evidence from Swiss-Prot and UniProt, consensus gene models were generated using EvidenceModeler and PASA. The final functional annotation integrated data from UniProt and InterProScan databases.

**Figure S3: Dotplot comparing the sequences of the Geisha subgenome assemblies:** Pseudomolecule comparison of the Geisha C subgenome with the Geisha E subgenome highlighting the dissimilarities that exist between the genomes from the respective progenitors. Orientation of the local alignments is color-coded, regions matching in both sequence and orientation are reported in blue while regions matching with inverted orientation are reported in orange.

**Figure S4: Dotplot comparison of the genome assemblies of Geisha and Red Bourbon varieties.** The diagram shows the high similarity that exists between the pseudomolecules of the two varieties. Orientation of the local alignments is color-coded, regions matching in both sequence and orientation are reported in blue while regions matching with inverted orientation are reported in orange.

**Figure S5: Distribution of sequence identity between *Coffea* assemblies.** *C. arabica* Geisha was binned into 25 Kbp windows and compared to the other *Coffea* genome assemblies. Each plot reports, for one pairwise comparison, the unscaled density distribution of the sequence identity measured for each window. **A)** *C. arabica* Geisha subgenomes C and E vs. with *C. Arabica* Red Bourbon subgenomes C and E, **B)** *C. arabica* Geisha subgenomes C and E vs. *C. canephora* and *C. eugenioides* genomes, **C) (left graphs)** *C. arabica* Geisha subgenome C vs *C. arabica* Geisha subgenomes E and *C. arabica* Red Bourbon subgenomes E, **C) (right graphs)** *C. arabica* Geisha subgenome E vs *C. arabica* Geisha subgenomes C and *C. arabica* Red Bourbon subgenomes C.

## References

Alonge M, Soyk S, Ramakrishnan S, Wang X, Goodwin S, et al. 2019. RaGOO: fast and accurate reference-guided scaffolding of draft genomes. Genome Biol. 20(1):224.

Altschul SF, Gish W, Miller W, Myers EW, Lipman DJ. 1990. Basic local alignment search tool. J Mol Biol. 215(3):403–10.

Blanco-Ulate B, Vincenti E, Powell ALT, Cantu D. 2013. Tomato transcriptome and mutant analyses suggest a role for plant stress hormones in the interaction between fruit and Botrytis cinerea. Front Plant Sci. 4:142.

Boot, W, 2013. Exploring the Holy Grail: Geisha coffee 10 years on. Roast Magazine May-June 2013.

Buchfink, Benjamin, Chao Xie, and Daniel H. Huson. 2014. ‘Fast and Sensitive Protein Alignment Using DIAMOND’. Nature Methods 12 (1): 59–60.

Carvalho A, 1985. Principles and practice of coffea plant breeding for productivity and quality factor: Coffea arabica. In: Coffee, Volume 4: Agronomy, Ed. RJ Clark and R Macrae, Elsevier Applied Science, London and New York 1985.

Chin, CS, Peluso P, Sedlazeck FJ, Nattestad M, Concepcion GT, et al. 2016. ‘Phased Diploid Genome Assembly with Single-Molecule Real-Time Sequencing’. Nature Methods 13(12): 1050–54.

Clarindo, WR, Carvalho, CR, Caixeta, E T, Koehler, AD 2013. Following the track of ‘Híbrido de Timor’ origin by cytogenetic and flow cytometry approaches. Genet. Resour. Crop Evol. 60:2253–2259.

Cros J, Combes MC, Chabrillange N, Duperray C, Angles AM, Hamon S 1995. Nuclear DNA content in the subgenus Coffea (Rubiaceae): inter- and intra-specific variation in African species. Can J Bot 73:14–20.

Cros J, Gavalda MC, Chabrillange N, Recalt C, Duperray C, Hamon S 1994. Variations in the total nuclear DNA content in African Coffea species (Rubiaceae). Café Cacao Thé 38:3–10.

DaMatta FM, Ronchi CP, Maestri M, Barros RS 2007. Ecophysiology of coffee growth and production, Braz. J. Plant Physiol., 19(4):485–510.

Denoeud F, Carretero-Paulet L, Dereeper A, Droc G, Guyot R, et al. 2014 The coffee genome provides insight into the convergent evolution of caffeine biosynthesis. Science 345(6201):1181–4.

Dereeper A, Guyot R, Tranchant-Dubreuil C, Anthony F, Argout X, et al. 2013. BAC-end sequences analysis provides first insights into coffee (*Coffea canephora* P.) genome composition and evolution. Plant Mol Biol. 83(3):177–89.

Dolezel J, Bartos J, Voglmayr H, Greilhuber J 2003. Nuclear DNA content and genome size of trout and human. Cytometry 51A: 127–128.

Gotz S, Garcia-Gomez JM, Terol J, Williams TD, Nagaraj SH, et al. 2008. High-throughput functional annotation and data mining with the Blast2GO suite. Nucleic Acids Res. 36:3420–3435.

Haas BJ. 2003. Improving the Arabidopsis genome annotation using maximal transcript alignment assemblies. Nucleic Acids Res. 31:5654–5666.

Haas BJ, Salzberg SL, Zhu W, Pertea M, Allen JE, et al. 2008. Automated eukaryotic gene structure annotation using EVidenceModeler and the Program to Assemble Spliced Alignments. Genome Biol. 9:R7.

ICO 2023, International Coffee Organization (ICO). Coffee report and outlook December 2023. (https://icocoffee.org/) (Accessed January 13, 20224).

Jones P, Binns D, Chang H-Y, Fraser M, Li W, et al. 2014. InterProScan5: genome-scale protein function classification. Bioinformatics 30:1236–1240.

Kent, W. J. 2002. BLAT—The BLAST-Like Alignment Tool. Genome Research 12 (4): 656–64.

Kim D, Langmead B, Salzberg SL. 2015. HISAT: a fast spliced aligner with low memory requirements. Nat Methods 12(4):357–60.

Koren S, Walenz BP, Berlin K, Miller JR, Phillippy AM. 2017. Canu: scalable and accurate long-read assembly via adaptive k-mer weighting and repeat separation. Genome Research. 27(5):722–736.

Krishnan, S. 2014. Genetic characterization of Geisha coffee. Conference paper, Denver Botanical Garden. Research Gate https://www.researchgate.net/publication/267358088e

Krug, CA, Mendes JET, Carvalho A 1949. Taxonomia de *Coffea arabica* L. II. *Coffea arabica* L. var Caturra e sua forma xanthocarpa. Bragantia 9(9-12) Campinas Set-Dez de 1949.

Kurtz S, Phillippy A, Delcher AL, Smoot M, Shumway M, et al.2004. Versatile and open software for comparing large genomes. Genome Biol. 5(2):R12.

Lashermes P, Hueber Y, Combes MC, Severac D, Dereeper A. 2016. Inter-genomic DNA Exchanges and Homeologous Gene Silencing Shaped the Nascent Allopolyploid Coffee Genome (Coffea arabica L.). G3 6(9):2937–48.

Li, Heng. 2018. ‘Minimap2: Pairwise Alignment for Nucleotide Sequences’. Edited by Inanc Birol. Bioinformatics 34 (18): 3094–3100.

Lomsadze A. 2005. Gene identification in novel eukaryotic genomes by self-training algorithm. Nucleic Acids Res. 33:6494–6506.

Montagnon C, Mahyoub A, Solano W, Sheibani F. 2021. Unveiling a unique genetic diversity of cultivated Coffea arabica L. in its main domestication center: Yemen. Genet Resour Crop Evol 68:2411–2422.

Morris, J. 2019. Coffee a global history. Reaktion Books Ltd., London 213 pp.

Quinlan, Aaron R., and Ira M. Hall. 2010. ‘BEDTools: A Flexible Suite of Utilities for Comparing Genomic Features’. Bioinformatics 26 (6): 841–42.

Sachs JD, Cordes KY, Rising J, Toledano P, Maennling N. 2019. Ensuring economic viability and sustainability of coffee production. Columbia Center on Sustainable Investment, https://papers.ssrn.com/sol3/papers.cfm?abstract_id=3660936

Salojärvi J, Rambani A, Yu Z, Guyot R, Strickler S, et al. 2024. The genome and population genomics of allopolyploid *Coffea arabica* reveal the diversification history of modern coffee Cultivars. Nat Genetics 56:721–731.

Scalabrin S, Magris G, Liva M, Vitulo N, Vidotto M, et al. 2024. A chromosome-scale assembly reveals chromosomal aberrations and exchanges generating genetic diversity in *Coffea arabica* germplasm. Nat Commun. 15(1):463.

Simão, F. A., Waterhouse RM, Ioannidis P, Kriventseva EV, and Zdobnov EM, 2015 BUSCO: assessing genome assembly and annotation completeness with single-copy orthologs. Bioinformatics 31: 3210–3212.

Slater G, Birney E. 2005. Automated generation of heuristics for biological sequence comparison. BMC Bioinformatics 6:31.

Smit A, Hubley R, Green P. 2013. RepeatMasker Open-4.0. http://www.repeatmasker.org 2013-2015.

Smit A, Hubley R. 2019. RepeatModeler-1.0. 11. Institute for Systems Biology. 2019. http://www.repeatmasker.org/RepeatModeler/.

Stanke M, Keller O, Gunduz I, Hayes A, Waack S, et al. 2006. AUGUSTUS: ab initio prediction of alternative transcripts. Nucleic Acids Res. 34:W435–W439.

Stoffel, K., van Leeuwen, H., Kozik, A., Caldwell, D., Ashrafi, H. et al. 2012. Development and application of a 6.5 million feature Affymetrix Genechip(R) for massively parallel discovery of single position polymorphisms in lettuce (Lactuca spp.). BMC Genomics 13: 185.

Vaser R, Sovic I, Nagarajan N, Sikic M “Fast and accurate de novo genome assembly from long uncorrected reads.“ Genome Res. 2017 May;27(5):737–746.

Walker BJ, Abeel T, Shea T, Priest M, Abouelliel A, et al. 2014. Pilon: an integrated tool for comprehensive microbial variant detection and genome assembly improvement. PLoS One. 2014 Nov 19;9(11):e112963.

Wang Y, Tang H, Debarry JD, Tan X, Li J, Wang X, et al. 2012. MCScanX: a toolkit for detection and evolutionary analysis of gene synteny and collinearity. Nucleic Acids Res. 40(7):e49.

WCR 2016 (World Coffee Research, Variety catalog, 2016, https://varieties.worldcoffeeresearch.org/varieties/geisha-panama)

